# Geography, not lifestyle, explains the population structure of free-living and host-associated deep-sea hydrothermal vent snail symbionts

**DOI:** 10.1101/2022.08.18.504305

**Authors:** Michelle A. Hauer, Corinna Breusing, Elizabeth Trembath-Reichert, Julie A. Huber, Roxanne A. Beinart

**Affiliations:** University of Rhode Island, Graduate School of Oceanography, Narragansett, RI, USA; Arizona State University, School of Earth and Space Exploration, Tempe, AZ, USA; Woods Hole Oceanographic Institution, Department of Marine Chemistry and Geochemistry, Woods Hole, MA, USA

**Keywords:** symbiosis, hydrothermal vents, population genomics, *Alviniconcha*, Mariana Back-Arc, microbial biogeography

## Abstract

Marine symbioses are predominantly established through horizontal acquisition of microbial symbionts from the environment. However, genetic and functional comparisons of free-living populations of symbionts to their host-associated counterparts are sparse. Here, we assembled the first genomes of the chemoautotrophic gammaproteobacterial symbionts affiliated with the deep-sea snail *Alviniconcha hessleri* from two separate hydrothermal vent fields of the Mariana Back-Arc Basin. We used phylogenomic and population genomic methods to assess sequence and gene content variation between free-living and host-associated symbionts. Our phylogenomic analyses show that the free-living and host-associated symbionts of *A. hessleri* from both vent fields are populations of monophyletic strains from a single species. Furthermore, genetic structure and gene content analyses indicate that these symbiont populations are differentiated by vent field rather than by lifestyle. Together, this work suggests that, despite the potential influence of host-mediated acquisition and release processes on horizontally transmitted symbionts, geographic isolation and/or adaptation to local habitat conditions are important determinants of symbiont population structure and intra-host composition.

## Background

Mutualistic animal-microbe associations are globally significant phenomena, shaping the ecology and evolution of both host animals and microbial symbionts (1). These symbiotic associations are maintained by transmission of symbionts from host parent to progeny either 1) directly, for example via the germline (vertical transmission), 2) indirectly, for example through an environmental population of symbionts (hereafter referred to as “free-living” symbionts) (horizontal transmission); or, 3) via a combination of both vertical and horizontal transmission (mixed mode transmission) (2).

Horizontal transmission is more commonly found in aquatic than terrestrial habitats, likely due to the ease with which microbes can be transported in water compared to air or soil (3). However, even for marine symbioses where horizontally transmitted microbial symbionts are observed in the environment (4), it is not yet clear whether free-living, environmental populations of symbionts represent host-associated populations at the strain level, or whether their diversity and composition differs. Free-living symbiont populations are likely shaped by the dynamic interactions with their host—for example, host animals may “seed” the environment by the release of their symbionts into the water column only upon host death (5) or via continuous release from live adults (6,7). In addition, ecological and evolutionary processes, such as dispersal barriers, natural selection, and genetic drift, can contribute to the diversity and biogeography of environmental symbionts (8,9).

Deep-sea hydrothermal vents are discontinuous, island-like habitats dominated by vent-endemic invertebrates that host primarily horizontally transmitted chemoautotrophic bacterial symbionts, making them opportune natural systems for understanding the biogeography of free-living microbial symbionts. In these mutualisms, the symbiotic bacteria are either obtained during a narrow competence window in early developmental stages or throughout the lifetime of the host (10,11) and are, in most cases, housed intracellularly within the host’s gill tissue. The symbionts oxidize chemical reductants (e.g., H_2_S, H_2_, CH_4_) in venting fluids to generate energy for the production of organic matter, thereby providing the primary food source for the host in an otherwise oligotrophic deep ocean (12) and accounting for the high ecosystem productivity characteristic of hydrothermal vents (13–15). Despite reliance on horizontal transmission, the majority of host species from hydrothermal vents affiliate with only one or two specific endosymbiont phylotypes (i.e., species or genera based on 16S rRNA gene sequence similarity) (12), possibly as a means to reduce the acquisition of cheaters (16). While a significant number of studies have focused on the diversity, composition and structure of the host-associated symbiont populations (e.g., 10,17–22), their free-living, environmental stages remain poorly investigated, partly due to the difficulty of detecting low abundance free-living symbionts in environmental samples. As a consequence, few free-living symbiont studies exist. Most of these studies have so far relied on investigations of particular marker genes (4,23); only one used an -omics approach but was limited to a single metagenome (24).

A recent shotgun metagenomic study found putative free-living symbiont populations of the provannid snail *Alviniconcha hessleri* in low-temperature diffuse venting fluids at two distinct vent fields of the Mariana Back-Arc, Northwest Pacific (15.5–18°N), (25), providing a unique opportunity to compare free-living and host-associated stages of chemosynthetic symbionts at hydrothermal vents. *Alviniconcha hessleri* belongs to the dominant fauna at hydrothermal vents in the Mariana Back-Arc Basin, where it lives in nutritional endosymbiosis with one species of sulfur-oxidizing, environmentally acquired Gammaproteobacteria (26,27). As an endemic species to the Mariana region, *A. hessleri* is currently listed as “Vulnerable” on the International Union for Conservation of Nature Red List of Threatened Species (https://www.iucnredlist.org), highlighting the need to identify the factors that contribute to its limited biogeographic range, including the population structure of its obligate microbial symbiont.

In this study, we applied phylogenomic and population genomic methods to evaluate the evolutionary relationships as well as the genetic and functional variation of *Alviniconcha hessleri* symbionts based on lifestyle by comparing free-living and host-associated symbiont populations collected from the same habitats. In addition, we addressed the effect of geography by comparing populations of both host-associated and free-living symbionts at two vent fields, Illium and Hafa Adai, that are approximately 140 km apart and differ notably in their geochemistry: Hafa Adai is known to support black smokers that emit high temperature fluids, with high amounts of hydrogen sulfide (H_2_S), whereas Illium is characterized by diffuse flow habitats that are lower in temperature and concentration of H_2_S (25).

## Methods

### Host-associated symbiont collection, sequencing, and genome assemblies

Three *A. hessleri* specimens each were collected from snail beds at the Illium vent field (3582 m) and the Voodoo Crater-2 (VC2) location within the Hafa Adai vent field (3277 m) in the Mariana Back-Arc Basin (Figure 1) using the remotely operated vehicle *SuBastian* on board the R/V *Falkor* in 2016. Symbiont-bearing snail gill tissues were dissected and stored in RNALater™ (Thermo Fisher Scientific, Inc.) at -80ºC until DNA extraction with the Zymo Quick DNA 96 Plus and ZR-96 Clean-up kits (Zymo Research, Inc.). High-throughput Illumina 150-bp paired-end libraries for all six samples were prepared and sequenced by Novogene Corporation, Inc. (Beijing, China) with an average yield of 58 million total Illumina reads (Supplementary Table 1). In addition, one gill sample from each vent field—Hafa Adai 172 (VC2) and Illium 13—was selected for long-read Nanopore sequencing on 2–3 MinION flow cells (Oxford Nanopore Technologies, Oxford, UK) using the SQK-LSK109 ligation kit (Supplementary Table 1).

**Figure 1.**
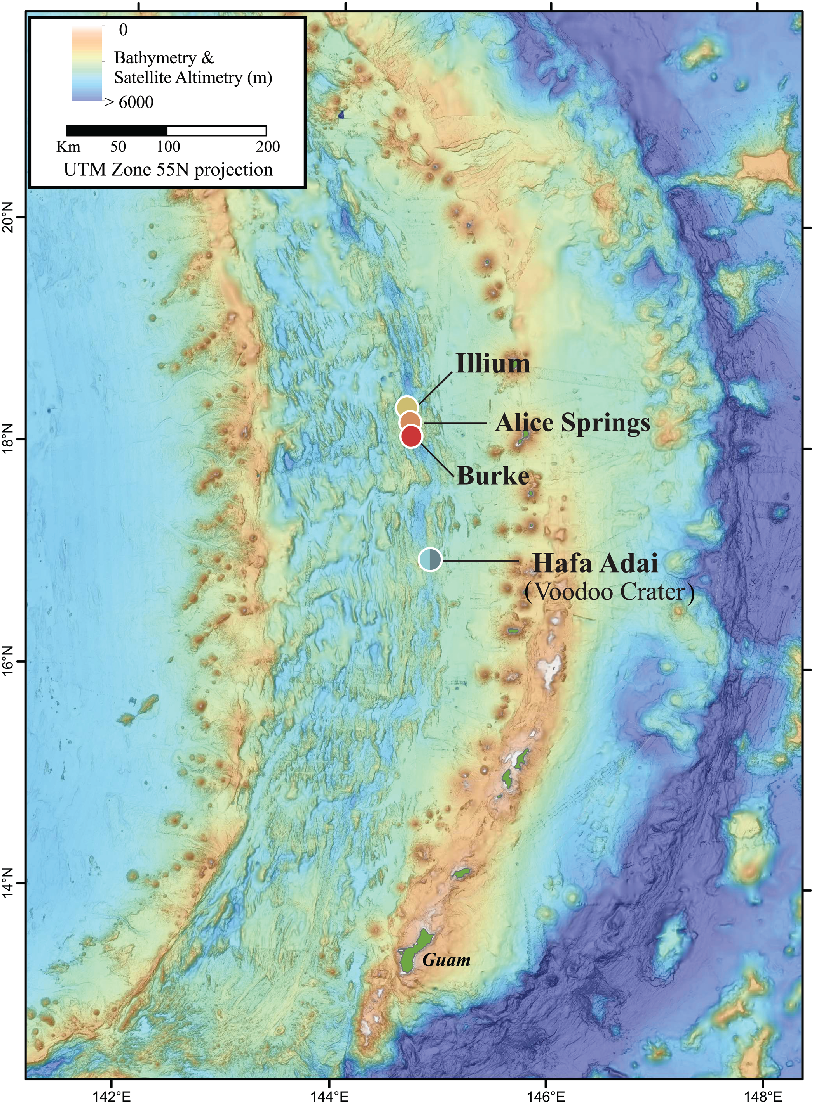
Map of the Mariana Back-Arc Basin adapted from ref. (25), indicating the locations of the two main hydrothermal vent fields sampled in this study, Illium and Hafa Adai. Free-living symbiont samples were obtained at low coverage from two additional vent sites investigated in ref. (25), Alice Springs and Burke. Colors associated with vent fields are used consistently across samples. A light cyan and teal are used for Hafa Adai, representing Voodoo Crater 1 and 2, respectively.

Raw Illumina reads were trimmed with Trimmomatic v0.36 (28) and common sequence contaminants were removed by mapping against the PhiX and human (GRCh38) reference genomes. Nanopore reads were base-called with Albacore (Oxford Nanopore Technologies, Oxford, UK) and adapter-clipped with Porechop (https://github.com/rrwick/Porechop). For Illium 13, symbiont Illumina and Nanopore reads were extracted through mapping against a draft co-assembly constructed with MEGAHIT (29) and reassembled with SPAdes v3.13.1 (30) using kmers between 21-91 in increments of 10. For Hafa Adai 172, all trimmed Illumina and Nanopore reads were assembled with metaSPAdes v3.13.1 (31) using the same parameters. Manual binning of the assemblies was conducted with GBTools (32) and contigs <200 bp and <500 bp were excluded from the Illium and Hafa Adai-VC2 assemblies, respectively. Assemblies were then scaffolded with SSPACE-Standard v3.0 (33) and SLR (34) and gapfilled with GapFiller v1-10 (35) and LR_GAPCLOSER (36). Final assemblies were polished with Pilon (37). Trimmed Illumina reads for the remaining samples—Illium 11, Illium 17, Hafa Adai 60, and Hafa Adai 64—were assembled with metaSPades v3.13.1 as described above and binned with MaxBin (38). All metagenome-assembled genomes (MAGs) were quality checked with Quast v5.0.2 (39) and CheckM v1.0.18 (40) and taxonomically assigned through the Genome Taxonomy Database toolkit (41).

### Free-living symbiont collection, sequencing, and genome assembly

High-quality MAGs of environmental *A. hessleri* symbionts from the Illium (GCA_003972985.1) and Hafa Adai-VC2 (GCA_003973075.1) vent sites were retrieved from a previous study (25). The fluid samples from which these MAGs were assembled were collected in direct vicinity of snail beds where the *A. hessleri* specimens were obtained (25). We further included a third MAG from diffuse-venting fluids at the Voodoo Crater-1 (VC1) (GCA_003973045.1) location within the Hafa Adai vent field, ∼5 meters from Hafa Adai-VC2. All details of the hydrothermal fluid collection, sample storage, sample processing, sequencing, assembly, and binning of metagenome-assembled genomes can be found in ref. (25). Information about raw sequencing reads is provided in Supplementary Table 1.

Raw Illumina metagenomics reads from the other vent sites sampled in ref. (25) —Burke, Alice Springs, Perseverance and Hafa Adai-Alba—were assembled and binned with the methods described above, but did not produce usable symbiont MAGs.

### Genome similarity and phylogenomic analyses

To confirm that all symbiont MAGs belong to the same bacterial species, we calculated average nucleotide identities (ANIs) via FastANI (42). A phylogenomic tree that included the six host-associated and the three free-living symbiont MAGs as well as reference genomes of other chemosynthetic Gammaproteobacteria (Supplementary Table 2) was then constructed with IQ-TREE2 (43) based on 70 single copy core genes in the Bacteria_71 collection (44). Parameter choice for phylogenomic reconstructions followed ref. (18).

### Population structure and gene content analysis

To determine symbiont population structure according to geography and lifestyle, we inferred DNA sequence polymorphisms in the free-living and host-associated samples by mapping metagenomic reads to a pangenome created with Panaroo (45) from all nine symbiont MAGs. Variants were called and filtered following the pipeline in ref. (18). All samples from Illium and Hafa Adai met our minimum 10x coverage threshold. Free-living samples from Burke and Alice Springs mapped at 5.9x and 3.9x coverage, respectively. The population structure and gene content analyses (see below) were repeated for these lower-coverage samples. All other free-living metagenomic samples from the remaining vent sites collected in ref. (25) had an insufficient number of reads mapped for further analyses. Principal coordinate analysis (PCoA) plots were created based on nucleotide counts converted to Bray-Curtis dissimilarities with the ggplot2 (46) and vegan (47) packages in Rstudio (48). To quantify the qualitative variant calling results depicted in the PCoAs, fixation indices (F_ST_) between individual samples were calculated following ref. (49) and plotted with pheatmap (50). The method from ref. (49) as well as scikit-allel (https://github.com/cggh/scikit-allel) were further used to calculate pairwise F_ST_ values between samples pooled by lifestyle or vent field.

Gene content variation among symbiont populations was determined via Pangenome-based Phylogenomic Analysis (PanPhlAn) (51) and visualized through a PCoA plot based on the Jaccard Similarity Coefficient. Genes that were uniquely associated with lifestyle and vent field, respectively, were extracted from the PanPhlAn gene presence/absence matrix. Functional predictions for these genes were either obtained from the Prokka (52) annotations created during pangenome construction or inferred by blasting the respective protein sequences against the NR database. Hypothetical and unknown proteins were further annotated via KEGG (53) and Alphafold (54). Differences in gene content between symbiont populations were visualized through Likert plots with the HH package (55) in RStudio.

### Validation of free-living symbiont populations

To gain confidence that the symbionts detected in our environmental samples represented truly “free-living” symbiont stages as opposed to symbionts associated with host larvae or shed gill cells, we calculated the ratio of symbiont 16S rRNA genes to host mitochondrial CO1 genes in all nine samples by mapping metagenomic reads to *Alviniconcha* symbiont 16S rRNA and host mtCO1 gene databases. To account for false positive mappings, we created additional background databases consisting of select bacterial (SUP05 clade bacteria, *Thiomicrospira*, and *Marinomonas*) and mollusk gene sequences. Bacterial 16S rRNA genes were downloaded from SILVA (56), while all *Alviniconcha* and mollusk mtCO1 genes were downloaded from BOLD (57). BBSplit (https://sourceforge.net/projects/bbmap/) was then used to separate *Alviniconcha* symbiont and host reads based on the taxon-specific and background 16S rRNA and CO1 gene databases.

## Results

### Free-living and host-associated symbionts belong to the same bacterial species

Our analysis included nine *A. hessleri* symbiont MAGs from the Illium and Hafa Adai vent fields: six host-associated symbiont genomes assembled in this study, and three previously published, free-living symbiont genomes from the diffuse venting fluids around *A. hessleri* beds (25) (Supplementary Table 3). All host-associated and two of the three free-living MAGs were of very high quality, with >90% completeness and <3% contamination. The third free-living MAG, Hafa Adai-VC1, had a medium quality (∼67% completeness). ANI values between all MAGs were >97.7% (Supplementary Table 4), suggesting that the nine *A. hessleri* symbiont genomes belong to the same bacterial species (42), *Thiolapillus* sp. based on the Genome Taxonomy Database, and confirming that the previously assembled free-living symbiont genomes were indeed *A. hessleri* symbionts (Supplementary Table 3). Corroborating the ANI results, the nine *A. hessleri* MAGs were monophyletic in our phylogenomic analysis relative to the gammaproteobacterial symbionts of other vent invertebrates (Figure 2). In agreement with phylogenetic analyses of the 16S rRNA gene (27), the nearest neighbors of the *A. hessleri* symbionts were the *Ifremeria nautilei* SOX symbiont, as well as *Thiolapillus brandeum*, a microbe not known to be symbiotic (58).

**Figure 2.**
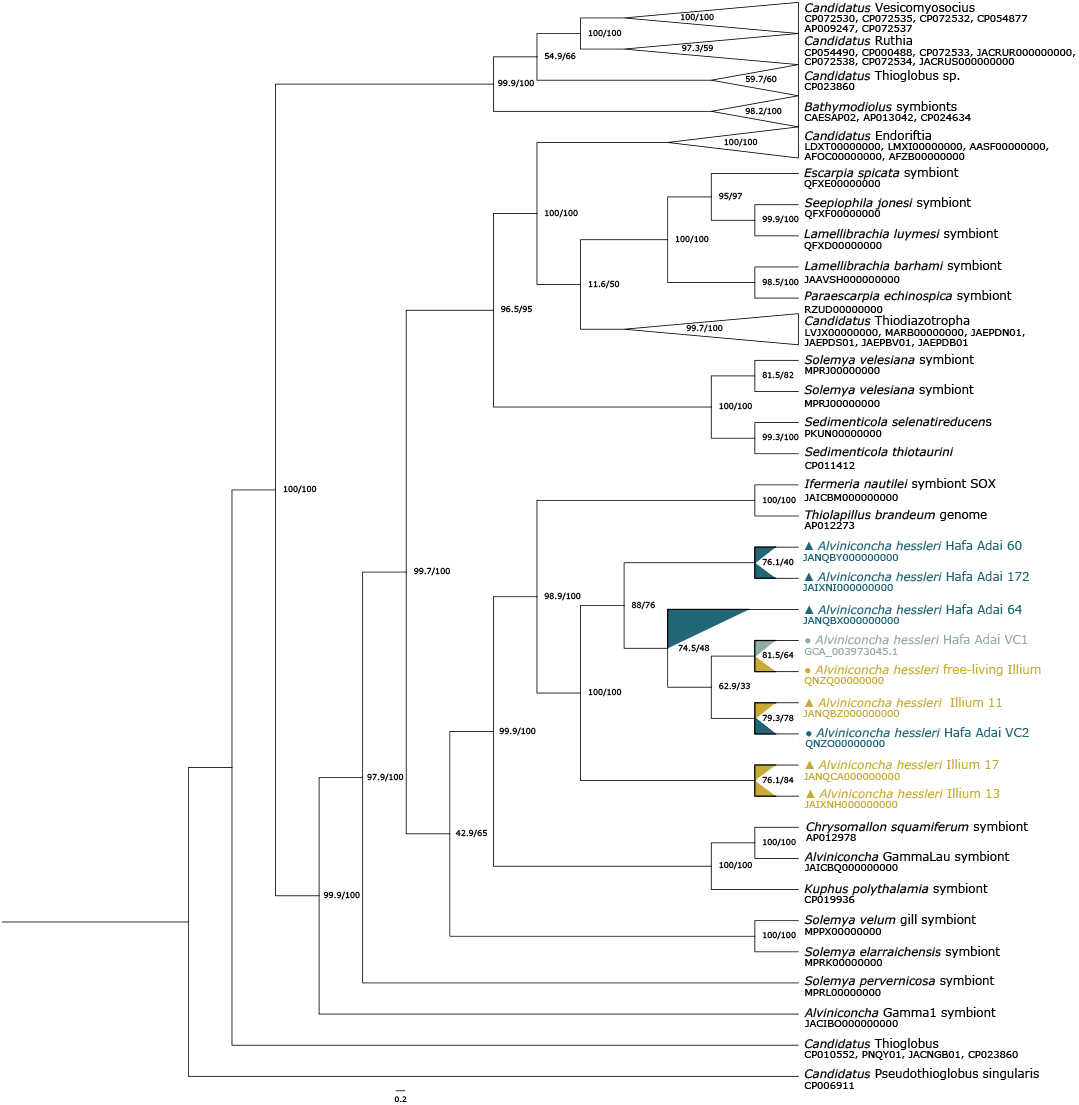
Cladogram branch transformed phylogenomic tree of chemosynthetic Gammaproteobacteria based on 70 single copy core genes. Genome accession numbers for all genomes included in the phylogenomic analysis can be found in Supplementary Table 1. *A. hessleri* symbionts are colored by vent field (Hafa Adai VC2: teal, Hafa Adai VC1: light cyan, Illium: yellow) with dots and triangles indicating free-living or host-associated lifestyle. *Candidatus* Pseudothioglobus singularis was used as outgroup for tree rooting.

### Environmental samples contain free-living symbiont populations

To investigate whether the symbionts observed in the diffuse flow samples were true free-living symbionts rather than symbionts associated with *A. hessleri* larvae or shed gill tissue, we calculated the ratio of symbiont 16S rRNA gene to host mitochondrial CO1 gene reads in all nine environmental and host-associated samples (Table 1). If the environmental symbiont samples were associated with larvae or host tissue debris/cells, we expect the ratio in the environmental and host-associated samples to be similar to one another. However, the 16S rRNA : mtCO1 ratio was consistently orders of magnitude higher in environmental samples than in host-associated samples, indicating a population of symbiont cells independent from host tissue. This finding provides evidence that our environmental samples include truly free-living *A. hessleri* symbiont populations.

**Table 1.** Percentage of unambiguous reads in environmental samples mapping to *Alviniconcha* symbiont 16S rRNA and mitochondrial CO1 genes, as well as the ratio of 16S rRNA to mtCO1.

| Sample | *Alviniconcha* 16S rRNA | *Alviniconcha* mtCO1 | 16S rRNA:CO1 | Origin |
| --- | --- | --- | --- | --- |
| Host-associated Hafa Adai 172 | 0.01859 | 0.00319 | 5.82758621 | Gill tissue |
| Host-associated Hafa Adai 60 | 0.01627 | 0.01112 | 1.4631295 | Gill tissue |
| Host-associated Hafa Adai 64 | 0.01706 | 0.01019 | 1.67419038 | Gill tissue |
| Host-associated Illium 11 | 0.01116 | 0.00844 | 1.32227488 | Gill tissue |
| Host-associated Illium 13 | 0.02004 | 0.00802 | 2.49875312 | Gill tissue |
| Host-associated Illium 17 | 0.01435 | 0.00721 | 1.99029126 | Gill tissue |
| Free-living Illium | 0.04302 | 0.00218 | 19.733945 | Diffuse fluids |
| Free-living Hafa Adai- VC2 | 0.08404 | 0.00193 | 43.5440415 | Diffuse fluids |
| Free-living Hafa Adai- VC1 | 0.06526 | 0.00027 | 241.703704 | Diffuse fluids |

### *A. hessleri* symbiont populations are structured primarily by vent field, not lifestyle

Our genome assemblies from both host tissue and diffuse vent fluids likely represent the dominant symbiont strain in each sample, but do not reveal the full extent of strain-level population variation between samples. To determine whether *A. hessleri* symbionts form subpopulations consistent with geography or lifestyle, we created a pangenome out of the individual symbiont MAGs that we used as reference for variant calling (Supplementary Tables 5, 6). Our variant detection method resulted in 2177 sequence polymorphisms for investigation of population genomic structure based on F_ST_ and ordination analyses (Figure 3).

**Figure 3.**
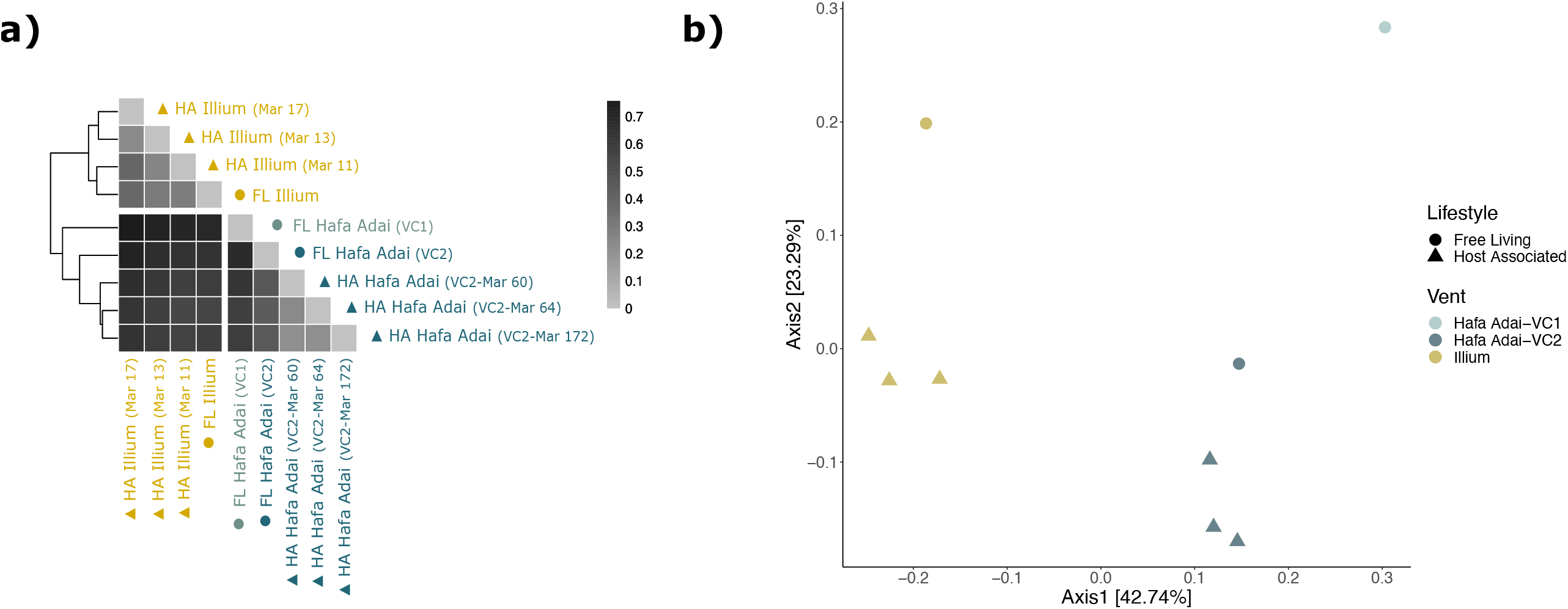
Symbiont population structure based on single nucleotide polymorphisms. A.) Heatmap of genome-wide, pairwise fixation indices (F_ST_) created using the pheatmap package in RStudio. F_ST_ values range from 0 to 1, where a value of 0 indicates no genetic differentiation, while a value of 1 indicates complete isolation among populations. “FL” and “HA” indicate free-living and host-associated symbionts, respectively. B.) PCoA plot based on Bray-Curtis distances illustrating the population structure of *A. hessleri* symbionts of different lifestyles and vent fields.

F_ST_ values were calculated pairwise between all nine populations (Figure 3a), as well as between samples pooled by lifestyle and vent field. F_ST_ values range from 0 to 1, where an F_ST_ value of 1 indicates that samples form genetically isolated subpopulations, while an F_ST_ value of 0 indicates that the samples form a single, well-mixed population. Between individual samples, pairwise F_ST_s showed a moderate (0.2–0.5) to strong (>0.5) differentiation, indicating that all samples represent distinct subpopulations with limited genetic exchange among each other (Figure 3a, Supplementary Table 7).

Genetic isolation among individual samples was typically stronger between (0.5395– 0.7560) than within (0.2123–0.4594) vent localities (i.e., Illium vs Hafa Adai-VC2). Within vent sites, the degree of differentiation was comparable among samples independent of lifestyle at Illium, while host-associated samples were more similar to one another than to free-living samples at Hafa Adai-VC2. When samples were pooled, overall pairwise F_ST_ values were markedly higher by vent field (0.4679 ± 0.0300 s.d.) than by lifestyle (0.0544 ± 0.0139 s.d.). The dominant effect of geography on symbiont population structure was supported by PCoAs where both free-living and host-associated samples from Illium clustered distinctly from Hafa Adai (VC1 and VC2) (Figure 3b). Despite the fact that Hafa Adai-VC1 and -VC2 differ spatially by only ∼5 meters, the free-living VC1 sample formed its own distinct subpopulation from both host-associated and free-living populations at VC2 (F_ST_: 0.5771–0.6731), suggesting very fine-scale geographic or environmental structuring.

These patterns were consistent in analyses based on 1271 and 793 variant sites that included the free-living, low coverage symbiont samples from Burke and Alice Springs, respectively (Supplementary Figure 1, 4; Supplementary Tables 8, 9). Burke represented the most divergent population, reaching F_ST_ values > 0.8 in all pairwise comparisons. Although Alice Springs clustered closely with free-living and host-associated symbionts from Illium in the PCoAs, F_ST_ values indicated a high degree of genetic isolation for this population (F_ST_ > 0.7). Analyses with samples pooled by vent field confirmed patterns of strong genetic differentiation between geographic locations without evidence for isolation-by-distance (Supplementary Table 10).

### *A. hessleri* symbiont gene content differs by vent field, not lifestyle

Gene content variation between symbiont populations was assessed based on lifestyle and geography. Similar to the population structure analyses, PCoA plots based on gene content variation across all nine host-associated and free-living populations revealed clustering by vent field but not by lifestyle: symbiont populations from Hafa Adai-VC1 and Hafa Adai-VC2 were more similar to one another than to Illium (Figure 4), although Hafa Adai-VC1 clustered as an independent population from all other samples.

**Figure 4.**
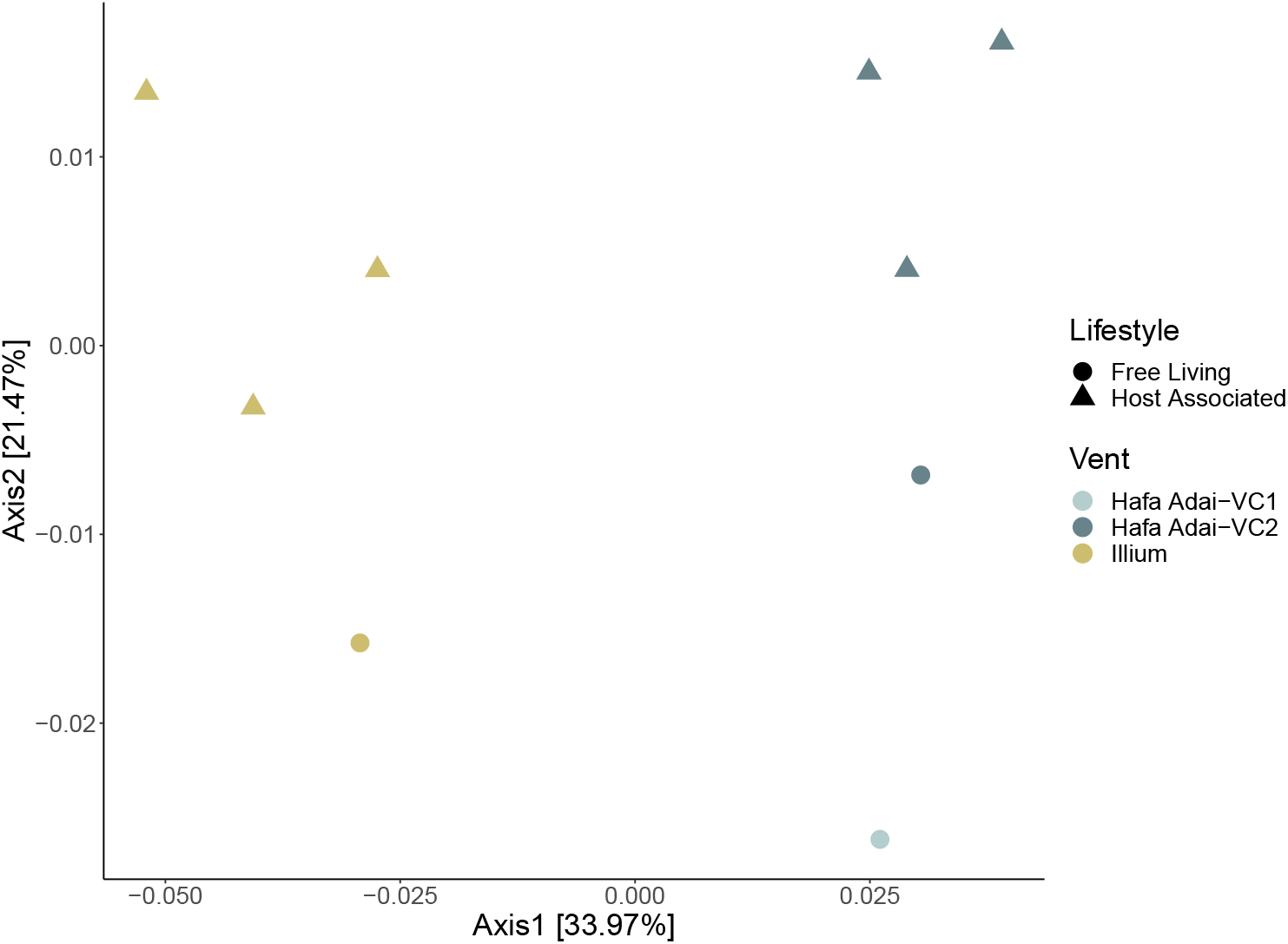
PCoA plot based on Jaccard distances illustrating the difference in gene content between *A. hessleri* symbionts based on both lifestyle and vent field.

Gene content differed more substantially by geography than by lifestyle: the Illium symbionts had 44 unique gene clusters, while the Hafa Adai (VC1 and VC2) symbionts had 26 (Figure 5a, Supplementary Table 11). Hypothetical and unknown proteins based on Prokka and the NR database were also assessed via KEGG and Alphafold, but yielded low confidence results. Of the successfully annotated genes unique to the Illium symbionts, most were predicted to be involved in the mobilome and DNA metabolism, followed by membrane transport, virulence, disease, defense; RNA metabolism; sulfur metabolism; cell signaling and regulation; conjugation; iron metabolism; glycolysis and gluconeogenesis; and detoxification and stress response. Genes unique to the Hafa Adai (VC1 & VC2) symbionts were predominantly associated with the mobilome, followed by membrane transport; RNA metabolism; motility and chemotaxis; DNA metabolism; virulence, disease and defense; and glycolysis and gluconeogenesis. Only three unique genes differentiated the host-associated and free-living symbiont populations across both vent fields (group_681, group_2104, group_2131, Supplementary Table 11). However, these genes could not be characterized by any database we used for functional annotations.

**Figure 5.**
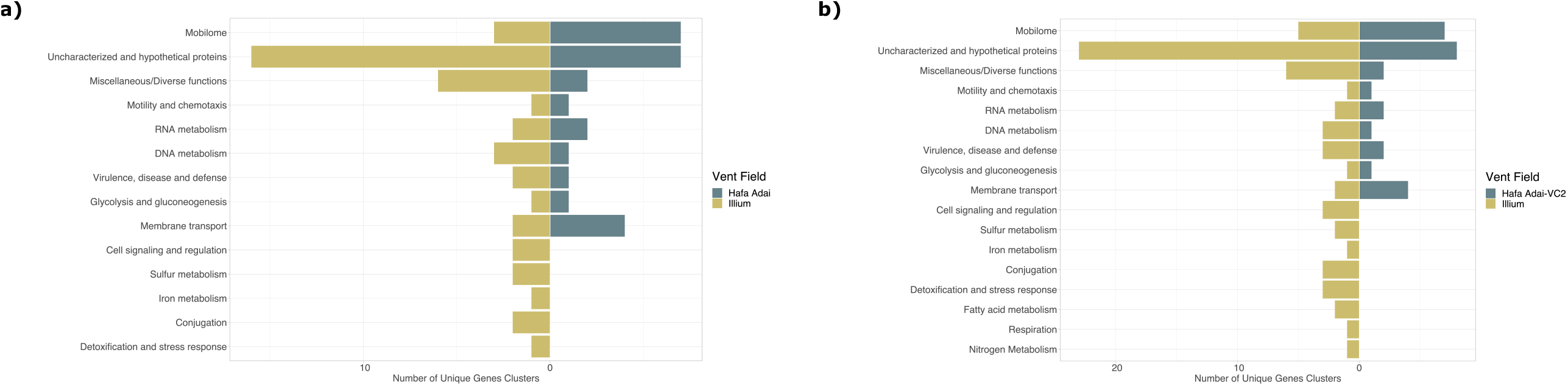
Likert plots showing A) number of unique genes between Illium and Hafa Adai (including both VC1 and VC2), and B) number of unique genes between Illium and Hafa Adai (VC2 only).

Given the small-scale geographic structuring found between VC1 and VC2 at Hafa Adai, we also compared the unique genes between Illium and Hafa-Adai VC2 symbionts alone (i.e., without VC1) (Supplementary Table 12, Figure 5b). In this case, there were an additional 18 unique genes for symbionts from Illium and 2 unique genes for symbionts from Hafa Adai-VC2 (Figure 5b). Only one of the genes unique to the Hafa Adai symbionts could be annotated and fell under the larger subcategory of “Virulence, Disease and Defense”, whereas unique genes of the Illium symbionts spanned a variety of metabolic functions.

Analyses that included symbiont reads from Alice Springs and Burke (Supplementary Figures 2, 3, 5; Supplementary Tables 13, 14) further supported the effect of geography over lifestyle on gene content variation in the *A. hessleri* symbionts. The population at Burke harbored a single unique, uncharacterized gene (Supplementary Table 13). When pooled with Illium as a “northern site”, additional genes unique to DNA metabolism and membrane transport were found, followed by genes involved in the mobilome, RNA metabolism, virulence, glycolysis and gluconeogenesis, cell signaling, conjugation and stress response (Supplementary Figure 3; Supplementary Table 13).

Alice Springs harbored three uncharacterized or hypothetical genes. When all three northern sites (Alice Springs, Illium, and Burke) were pooled together, seven unique genes were found. Four of these were related to DNA metabolism, virulence, conjugation and transposition (Supplementary Table 14). Since Alice Springs and Illium are more geochemically similar to one another than either vent is to Burke, we also investigated the unique genes shared by these two vent fields alone: four unique genes were found, one of which fell under the functional category of Virulence, Disease and Defense.

## Discussion

Here, we compared free-living and host-associated symbiont populations of *Alviniconcha hessleri* from two vent fields in the Mariana Back-Arc. Based on ANI and taxonomic assignments, our nine representative, medium- to high-quality MAGs can be considered to represent a single species within the genus *Thiolapillus* (58). Our results provide strong evidence that diffuse fluid flow microbial communities include populations of free-living symbionts, further supporting an expected model of horizontal transmission in *Alviniconcha* species (18,59).

Both population structure and gene content analyses suggest that *A. hessleri* symbionts form subpopulations that segregate by geography more strongly than by lifestyle. These patterns agree with previous studies of host-associated *A. hessleri* symbiont biogeography at the 16S rRNA gene level (27) and with observations in other horizontally-transmitted associations from hydrothermal vents, such as bathymodiolin mussels (49,60) and provannid snails (18), that have been shown to partner with habitat-specific symbiont strains. These results are also consistent with studies of non-symbiotic hydrothermal vent microbial communities, which show that microbes are shaped by their local environment (61). Uptake of environmental symbiont strains bears a risk of infection to the host by cheaters (16), but also enhances an animal’s ability to flexibly associate with locally available symbiont strains and, therefore, to maximize the habitat range in which they can settle (2,18). Hydrothermal vents are ephemeral and geochemically dynamic habitats, harboring microbial communities that are shaped by environmental conditions both within and between vent fields (61); therefore, it may be ecologically and evolutionarily advantageous for vent animals to acquire symbiont strains that are likely locally adapted (62).

The dynamics of microbial interaction with the host during acquisition and release processes can have significant impacts on the population structure and composition of horizontally transmitted symbionts. It is not known whether *A. hessleri* can replenish or recycle its symbionts, or if symbiont acquisition occurs only once upon settlement. For example, hydrothermal vent tubeworms seed the environment with their symbionts only upon death (5), *Bathymodiolus* mussels can acquire and release their symbionts throughout their lifetime (11,63), and *Vibrio fischeri* symbionts are expelled every morning by their sepiolid squid host (64). In *V. fischeri*, it is well established that evolution in the free-living stage—for example, via horizontal gene transfer—impacts the evolution of host-microbe interactions, though the role of novel mutations remains unclear (63). Although *A. hessleri* symbionts were overall more strongly partitioned by geography than by lifestyle, all symbiont samples were genetically distinct from each other and formed separate free-living or host-associated subpopulations. These findings suggest that symbiont exchanges between host and environment throughout the lifetime of the host are limited but might occur occasionally via symbiont uptake or release (49), thereby leading to mixing of host-associated and free-living symbiont pools. Periodic switching of symbiont strains could increase shared genetic variation among intra- and extra-host symbiont populations, while maintaining geographic differentiation in the presence of dispersal barriers and/or environmental selection. All samples from Illium showed a comparable degree of differentiation from each other, while samples from Hafa Adai were notably divergent between free-living and host-associated lifestyles. These patterns could arise from differences in the sampling locations of the free-living symbiont populations (e.g., distance from the snail beds) and/or the age of the *Alviniconcha* host individuals. Although we do not have size-related data for the collected specimens, it is possible that the snail individuals from Hafa Adai were older than those from Illium, giving host-associated symbiont populations more time to diverge from their free-living counterparts. Strong genetic differentiation between host-associated and free-living symbiont populations can be expected if hosts take up similar symbiont strains that have limited exchange with the environment post-infection, while the free-living symbiont population experiences more turnover.

The high genetic isolation of symbiont populations observed between vent fields may reflect the influence of both neutral (e.g., dispersal barriers and isolation by distance) and selective processes (e.g., adaptation to habitat differences between vent fields) on symbiont biogeography. Illium, Burke, and Alice Springs are all northern vent fields within the Mariana Back-Arc Basin that are characterized by sites of low-temperature diffuse fluid flow, while Hafa Adai is located further south and contains high-temperature black smokers (25). Illium and Alice Springs are similar geochemically, notably in that they are both low in H_2_S concentrations. The close proximity (∼360 m) and overlap in geochemical characteristics between Alice Springs and Illium may explain why these vent fields clustered together in our population structure analyses. By contrast, Burke is located about 4 km from both Illium and Alice Springs and has a distinct geochemical signature, which could contribute to the high genetic isolation seen for this vent field. Overall, however, no clear pattern of isolation-by-distance was observed, indicating that ecological factors might play a more important role than dispersal barriers in shaping symbiont population structure, in agreement with the oceanographic connectivity between the northern and central Mariana Back-Arc Basin (65).

Interestingly, Hafa Adai-VC1 — while more similar to Hafa Adai-VC2 than any other vent site — represented its own symbiont sub-population, suggesting small scale population structuring of symbionts within vent fields. Local patchiness of symbionts, as observed in our study, mirrors patterns found for host-associated symbionts of cold-seep vestimentiferan tubeworms (66) and *Acropora* corals (67). Although Hafa Adai-VC1 and -VC2 were only ∼5 meters apart, it is possible that *Alviniconcha hessleri* symbionts have extremely low dispersal potential that could be further reduced by small-scale circulation within vent sites due to physical structuring in the subseafloor (68,69). Alternatively, micro-niche adaptation driven by locally fluctuating environmental conditions might contribute to these patterns.

Among the identified differences in gene content, symbionts from Illium uniquely harbored genes related to iron and sulfur metabolism. As iron concentrations appear to be reduced at northern Mariana Back-Arc vents (70) and are typically lower in diffuse flow than black smoker fluids, it is possible that symbionts at Illium harbor high affinity Fe^2+^ transporters to efficiently obtain this essential element for their metabolism. All symbiont populations, including the low coverage samples from Alice Springs and Burke, showed differences in the presence of genes related to the mobilome and virulence, disease and defense. This suggests that each vent field supports distinct viral communities, as hydrothermal vent viruses have restricted bacterial and archaeal host ranges, and viral communities are typically endemic to a given vent site due to limited dispersal or environmental selection (71,72). The high number of unique genes related to the mobilome may be a consequence of integrated phage-derived genetic material that reflect the local, free-living viral communities.

## Conclusions

Our research demonstrates that *Alviniconcha hessleri* symbiont populations are primarily structured by geography rather than by their host-associated or free-living lifestyle. Future work using population genomic approaches should help clarify the predominant force(s) shaping the geographic population structure, as recent analyses of the symbionts associated with other *Alviniconcha* species suggests that both genetic drift and local adaptation play a role in symbiont biogeography (18). Although our analyses indicate a weak effect of lifestyle on symbiont genetic structure, it is possible that free-living and host-associated populations are characterized by differences in gene expression. A comparison of gene expression between lifestyles may provide additional clarity on the extent to which these symbiont sub-populations differ functionally. Our work also strengthens previous evidence for horizontal symbiont transmission in *Alviniconcha* species (18,59), despite the fact that almost nothing is currently known about the dynamics of symbiont acquisition and release in these species. Given that *A. hessleri* has been classified as “Vulnerable” on the IUCN Red List (https://www.iucnredlist.org) and is a dominant species at vents that are part of the Marianas Trench Marine National Monument, it is critical for future conservation and management that we understand the genetic connectivity of the symbiotic microbes that support this foundation species.

## Declarations

### Data availability

The datasets supporting the conclusions of this article are available in the National Center for Biotechnology Information repository, under BioProject number PRJNA763533. The previously published, free-living raw sequencing reads and corresponding MAGs are available at the National Center for Biotechnology Information under BioProject PRJNA454888.

## Acknowledgements

We thank the Schmidt Ocean Institute, the captain, crew and pilots of the R/V *Falkor* and ROV *SuBastian*, as well as Bill Chadwick, David Butterfield, Verena Tunnicliffe, and Amanda Bates for their support in the sample collections. The data collected in this study includes work supported by the Schmidt Ocean Institute during the R/V *Falkor* cruise FK161129. This work was funded by the National Science Foundation (grant number OCE-1736932 to RAB and Graduate Research Fellowship to MAH). JAH was funded by the NOAA Ocean Exploration and Research (OER) Program and the NSF Center for Dark Energy Biosphere Investigations (C-DEBI) (OCE-0939564). ETR was supported by the NASA Postdoctoral Fellowship with the NASA Astrobiology Institute and the L’Oréal USA For Women in Science Fellowship.

