## Supplementary figures and images for "Geography, not lifestyle, explains the population structure of free-living and host-associated deep-sea hydrothermal vent snail symbionts"

### Supplemental Figure 1

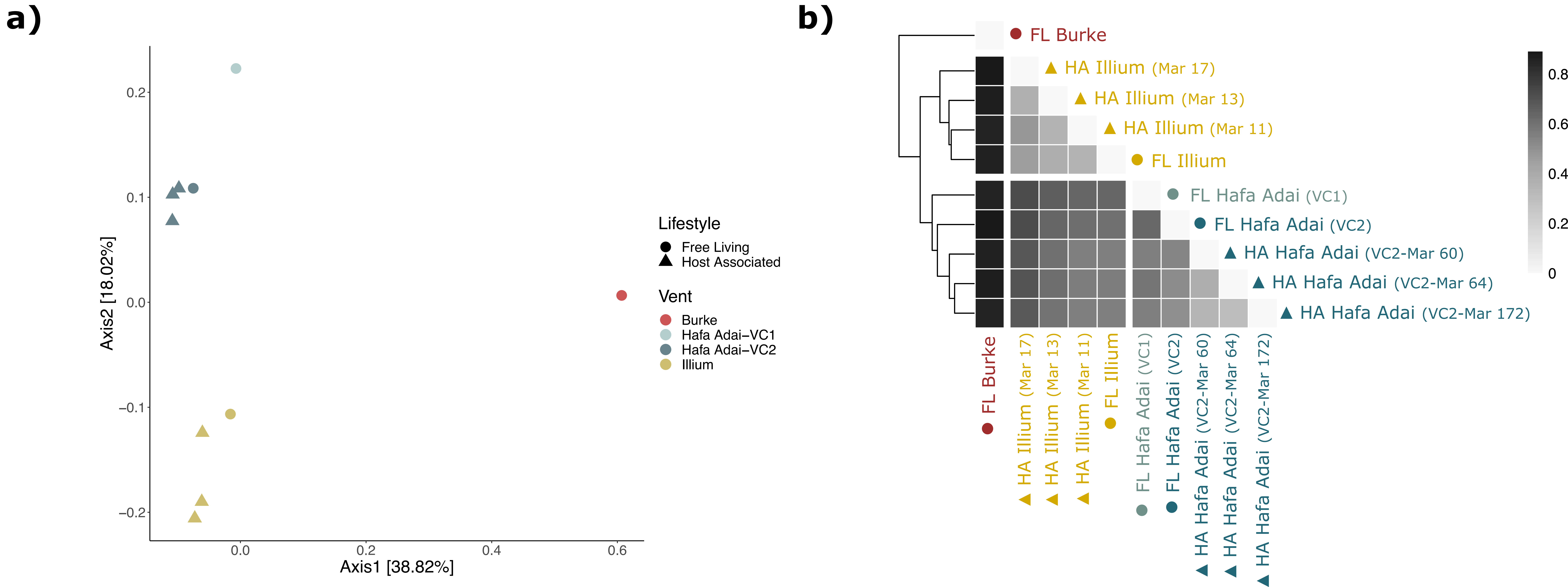

### Supplemental Figure 2

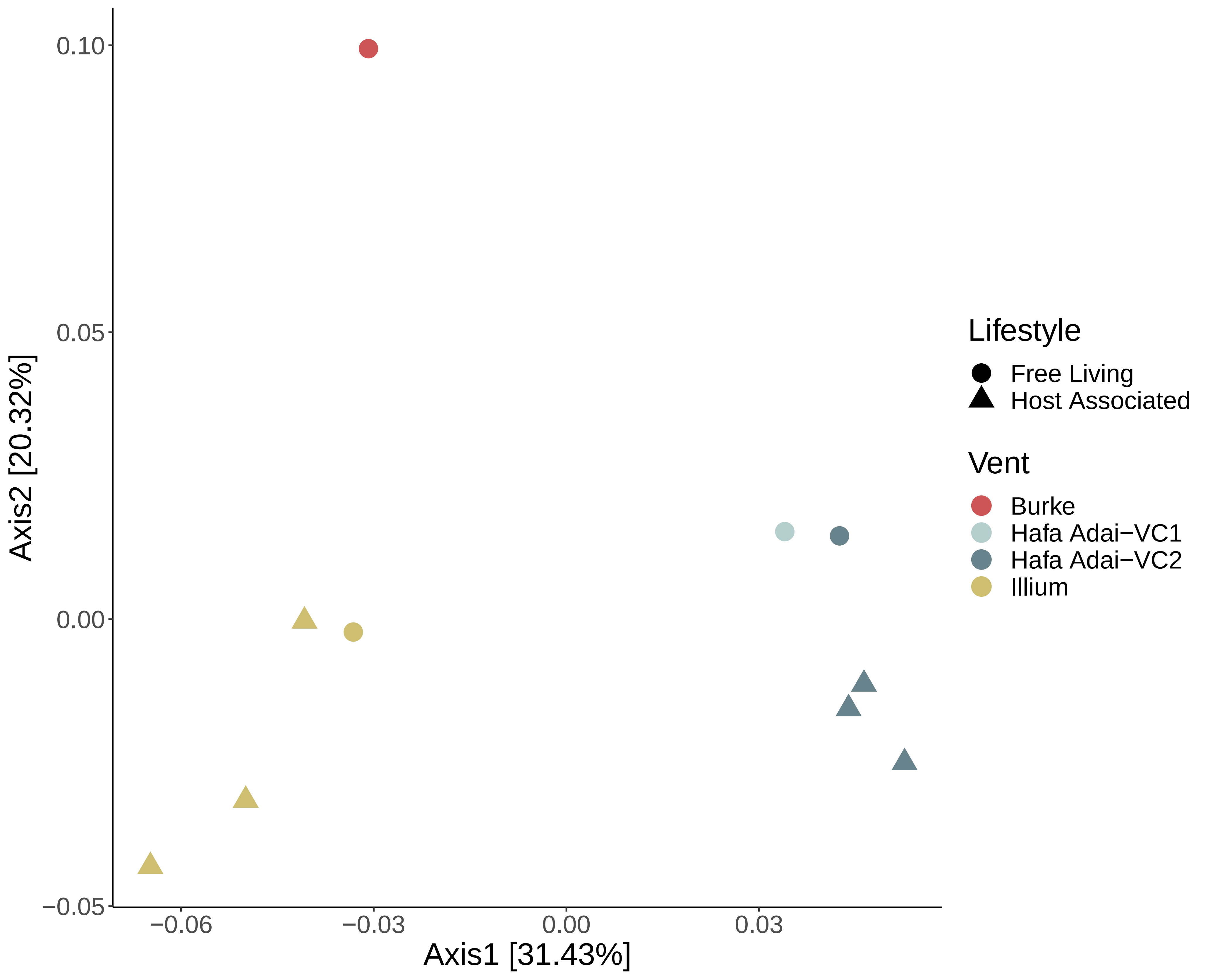

### Supplemental Figure 3

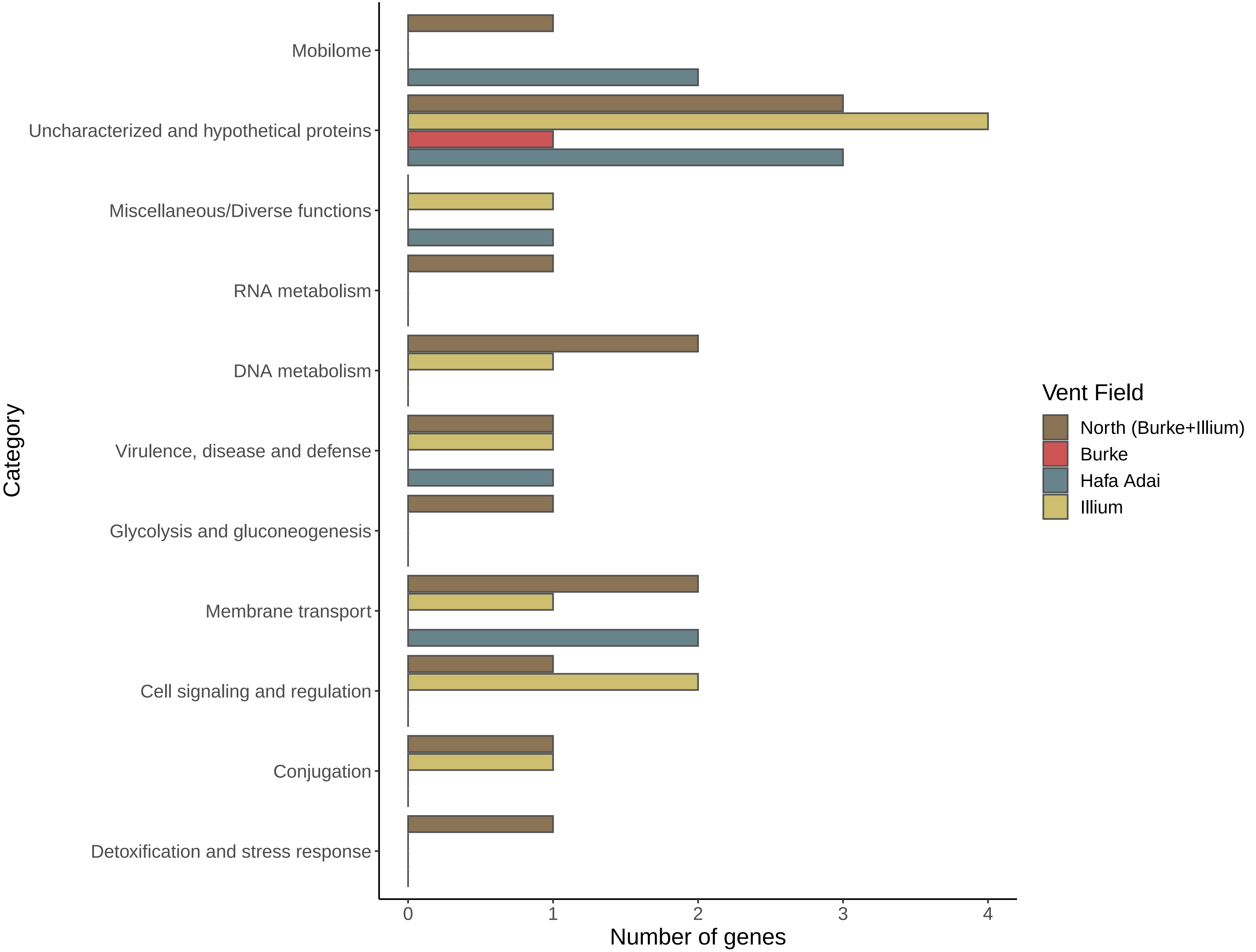

### Supplemental Figure 4

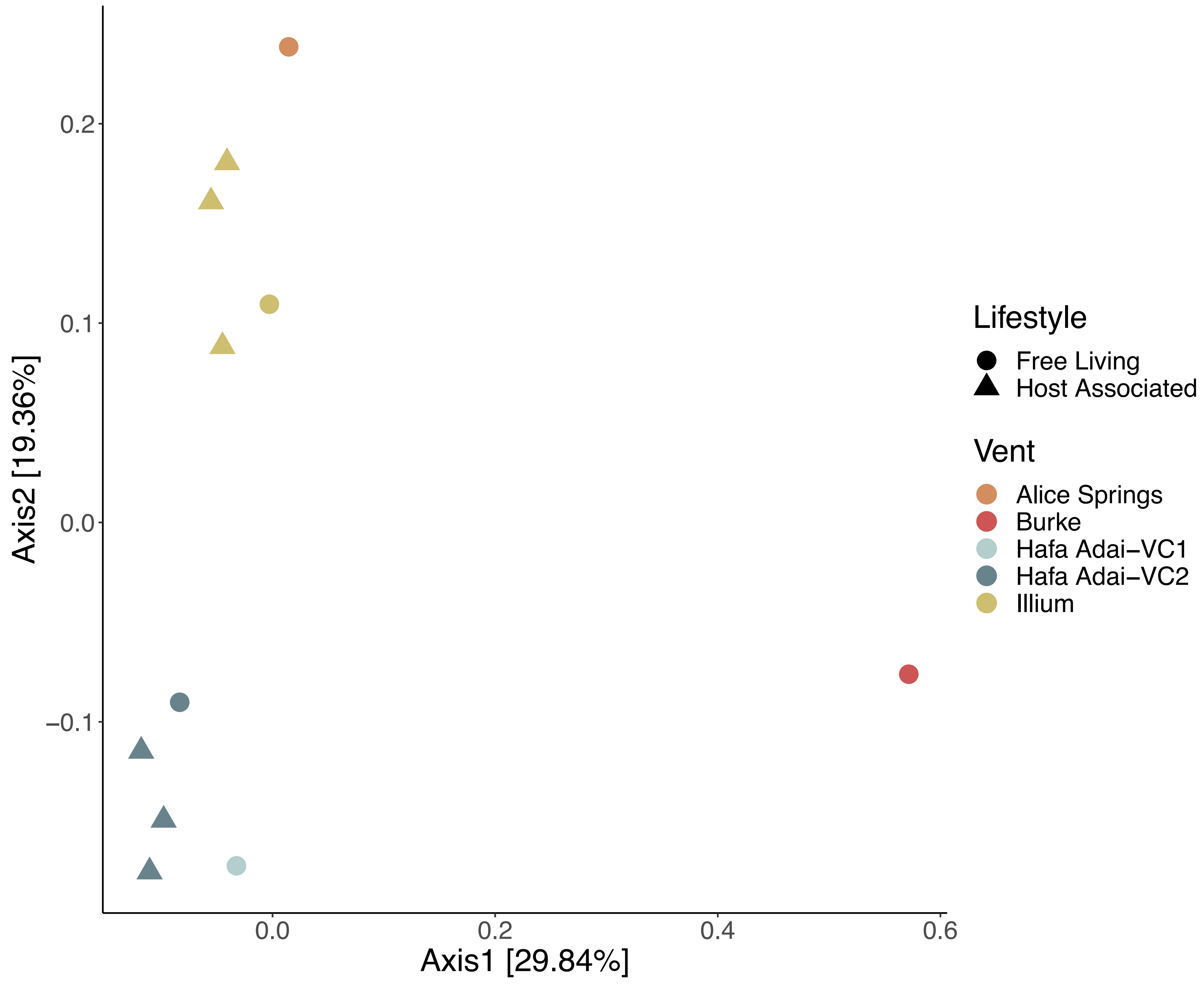

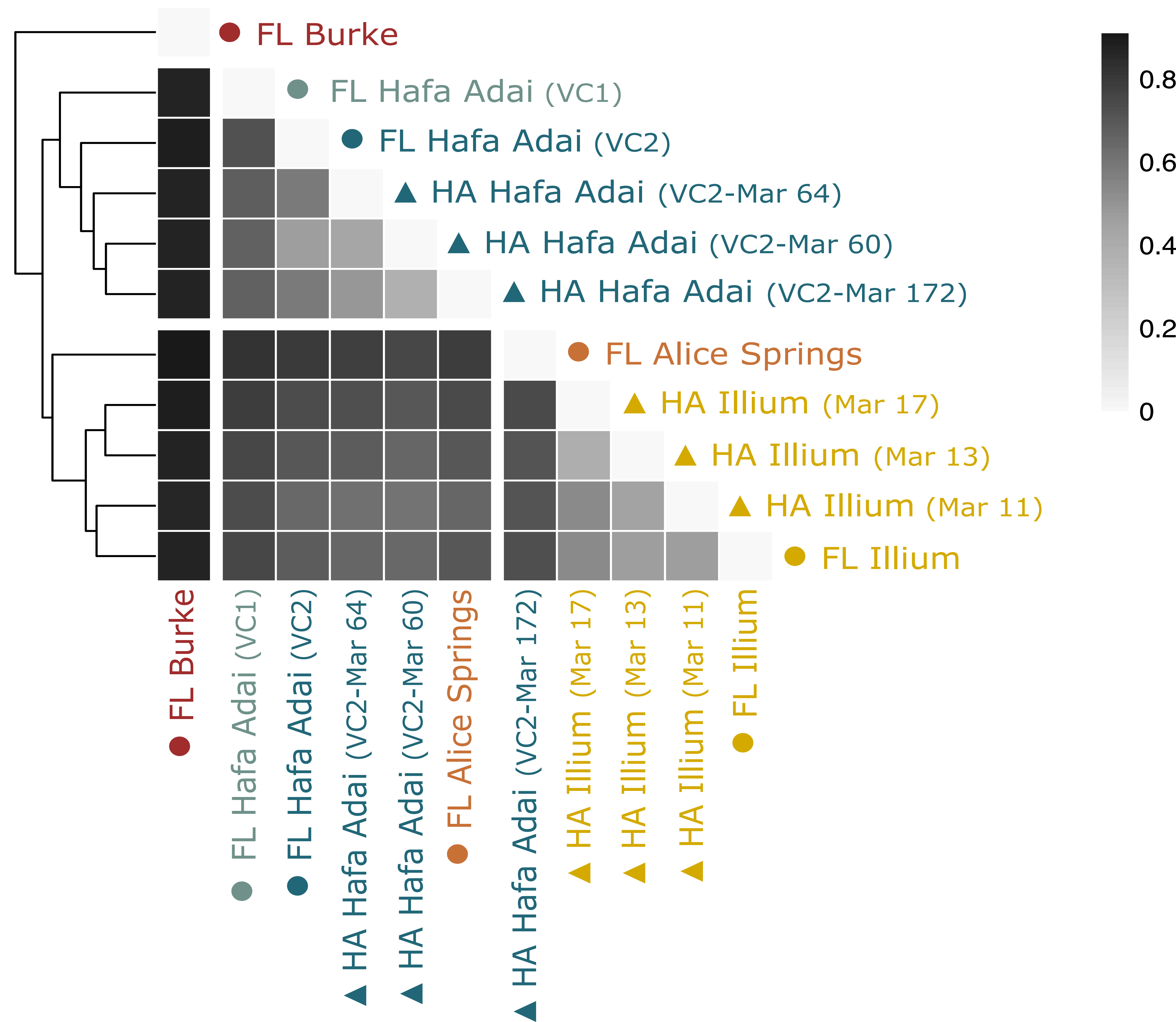

### Supplemental Figure 5

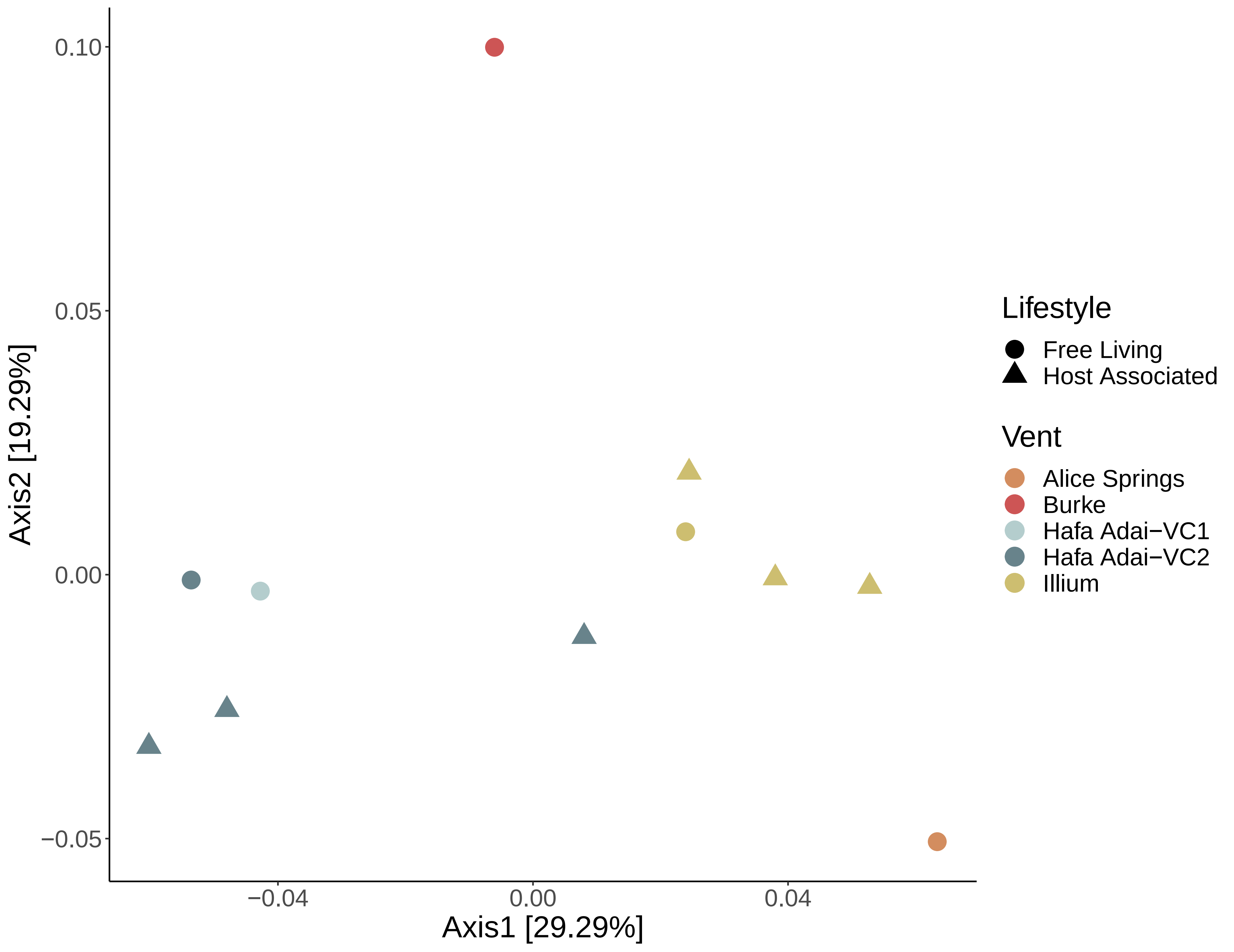
