## Supplemental Figure Captions for "Geography, not lifestyle, explains the population structure of free-living and host-associated deep-sea hydrothermal vent snail symbionts"

**Supplementary Figure Captions**

**Supplementary Figure 1.** Population structure analyses based on 1271 single nucleotide polymorphisms. Includes all nine samples from Illium and Hafa Adai plus symbiont reads from Burke. A) PCoA plot based on Bray-Curtis distances exhibiting population genetic structuring by geography and lifestyle. B) Heatmap of genome-wide, pairwise fixation indices (F_ST_) created using the pheatmap package in RStudio. F_ST_ values range from 0 to 1, where a value of 0 indicates no genetic differentiation, while a value of 1 indicates complete isolation among populations. “FL” and “HA” indicate free-living and host-associated symbionts, respectively.

**Supplementary Figure 2.** PCoA plot based on Jaccard distances illustrating the difference in gene content between *A. hessleri* symbionts based on both lifestyle and vent field, including all nine samples from Illium and Hafa Adai in addition to the free-living symbiont population from the Burke vent field.

**Supplementary Figure 3.** Barplot showing the number of unique gene clusters per functional category between vent fields, including a “northern” site category which combines the Illium and Burke vent fields. Created using ggplot in RStudio.

**Supplementary Figure 4.** Population structure analyses based on 793 single nucleotide polymorphisms. Includes all nine samples from Illium and Hafa Adai, plus symbiont reads from both the Burke and Alice Springs vent fields. A) PCoA plot based on Bray-Curtis distances exhibiting population genetic structuring by geography and lifestyle. B) Heatmap of genome-wide, pairwise fixation indices (F_ST_) created using the pheatmap package in RStudio. F_ST_ values range from 0 to 1, where a value of 0 indicates no genetic differentiation, while a value of 1 indicates complete isolation among populations. “FL” and “HA” indicate free-living and host-associated symbionts, respectively.

**Supplementary Figure 5.** PCoA plot based on Jaccard distances illustrating the difference in gene content between *A. hessleri* symbionts based on both lifestyle and vent field, including all nine samples from Illium and Hafa Adai in addition to free-living symbiont samples from the Burke and Alice Springs vent fields.
